# Human fecal microbiota is associated with colorectal cancer

**DOI:** 10.1101/2020.11.17.387647

**Authors:** Qiulin Yao, Meifang Tang, Liuhong Zeng, Zhonghua Chu, Hui Sheng, Yuyu Zhang, Yuan Zhou, Hongyun Zhang, Huayan Jiang, Mingzhi Ye

## Abstract

**Background:** Colorectal cancer (CRC) is one of the most common cancers. In recent studies, the gut microbiota has been reported to be potentially involved in aggravating or favoring CRC development. However, little is known about the microbiota composition in CRC patients after treatment. In this study, we explored the fecal microbiota composition to obtain a periscopic view of gut microbial communities. We analyzed microbial 16S rRNA genes from 107 fecal samples of Chinese individuals from three groups, including 33 healthy individuals (Normal), 38 CRC patients (Fa), and 36 CRC post-surgery patients (Fb).

**Results:** Species richness and diversity were decreased in the Fa and Fb groups compared with that of the Normal group. Partial least squares discrimination analysis showed clustering of samples according to disease with an obvious separation between the Fa and Normal, and Fb and Normal groups, as well as a partial separation between the Fa and Fb groups. Based on linear discriminant analysis effect size analysis and a receiver operating characteristic model, *Fusobacterium* was suggested as a potential biomarker for CRC screening. Additionally, we found that surgery greatly reduced the bacterial diversity of microbiota in CRC patients. Some commensal beneficial bacteria of the intestinal canal, such as *Faecalibacterium* and *Prevotella*, were decreased, whereas the drug-resistant *Enterococcus* was visibly increased in CRC post-surgery group. Meanwhile, we observed a declining tendency of *Fusobacterium* in the majority of follow-up CRC patients who were still alive approximately 3 y after surgery. We also observed that beneficial bacteria dramatically decreased in CRC patients that recidivated or died after surgery. This revealed that important bacteria might be associated with prognosis.

**Conclusions:** The fecal bacterial diversity was diminished in CRC patients compared with that in healthy individuals. Enrichment and depletion of several bacterial strains associated with carcinomas and inflammation were detected in CRC samples. *Fusobacterium* might be a potential biomarker for early screening of CRC in Chinese or Asian populations. In summary, this study indicated that fecal microbiome-based approaches could be a feasible method for detecting CRC and monitoring prognosis post-surgery.

## 1. Introduction

Colorectal cancer (CRC) is the third most common cancer worldwide, with the annual occurrence of 1 360 000 new cases and more than 600 000 deaths^1,2^. Because of its high incidence, increased difficulty in early diagnosis, and high mortality rate, colorectal cancer has become a major public health issue, especially in less developed regions. Moreover, survival and risk of recurrence have been reported to vary based on the stage of the tumor. According to the pathological classification, in cases of tumors confined to stages I and II, resection surgery can be curative with a 5-y survival rate of up to 80 %; however, the prognosis is dramatically decreased in tumors at a later stage, due to the increased occurrence of metastasis^3^. Therefore, a new diagnostic method for the early detection of lesions that would be noninvasive and easy to perform, is gaining attention among researchers.

Gut microbiota have been suggested to be potentially involved in the development of colorectal cancer. Bacteria and their related products might participate in the initiation or progression of sporadic colorectal cancer by a variety of mechanisms, including induction of inflammation, production of mutagenic toxins and reactive oxygen species (ROS), and the conversion of pro-carcinogenic dietary factors into carcinogens. These mechanisms have been shown to result in DNA and RNA damage, directly or indirectly inhibit DNA repair^1,4,5^, affect specific signal pathways, and block antitumor immunity^2^. Several bacteria have been reported to exhibit a carcinogenic risk. *Escherichia coli, Streptococcus bovis*, and *Bacteroides fragilis* are the bacteria most often described to be associated with colonic neoplasia^6^. The colibactin-producing *E. coli* has been reported to directly attack the host DNA, by introducing DNA breaks that lead to genomic instability and increased mutation frequency^4,7^. Whereas, *Enterococcus faecalis* is known to indirectly lead to DNA damage in the epithelium by inducing high levels of ROS^8,9^, which are typically produced by host cells during inflammation. *Fusobacterium nucleatum* has been reported to modulate the tumor-immune microenvironment, potentiating intestinal tumorigenesis in mice^10^. In addition, some studies have indicated that *F. nucleatum* is enriched in the gut of CRC patients^11,12^ and have even suggested it as a putative prognostic factor in CRC^13,14,15^.

Changes in the abundance of some gut commensal bacteria have been linked to dysbiosis observed in several human diseases. One such case regards *Faecalibacterium prausnitzii*, a protective bacterium, which was found to be decreased in CRC patients^6^. Culture supernatants of *F. prausnitzii* were shown to protect mice against 2,4,6-trinitrobenzenesulfonic acid-induced colitis, a potent risk factor for colon cancer^16^. The collection of fecal samples, in which the microorganism composition is known to be highly correlated with the colonic lumen and mucosa, seems to be an ideal approach, as this data can provide a periscopic view of gut microbial communities^17^, without the need for invasive procedures, such as colonoscopies.

To date, a large amount of research has focused on the gut microbiota of CRC patients; however, the microenvironmental changes in the colorectum of patients after therapy, such as surgery, chemotherapy, or radiotherapy, have not been widely studied. As specific bacteria might drive tumorigenesis, we aimed to identify whether the population of these bacteria were decreased after effective treatment. If so, it would indicate that effective treatment might result in the alteration of the microbiota of CRC patients to one more similar to that of normal samples.

Thus, to understand the structure of the gut microbial community and the changes post-surgery in CRC patients, we investigated the microbiota in the stools of CRC patients, CRC patients after surgery, and healthy individuals using 16S rDNA amplicon sequencing.

## 2. Results

### Summary of the study

Our study population was composed of 33 healthy individuals (Normal), 38 CRC patients before treatment (Fa), and 36 CRC patients after surgery (Fb) **(Table 1**).

**Table 1.** Demographic structure and clinical data of the study population

|  | **Fa** | **Fb** | **Normal** |
| --- | --- | --- | --- |
| **Number** | 38 | 36 | 33 |
| **Male, n (%)** | 24 (63.16) | 23 (63.89) | 17 (51.52) |
| **Mean age (±SD, y)** | 64.32 ± 11.14 | 63.19 ± 10.73 | 59.00 ± 4.94 |
| **Pathological stage** |  |  |  |
| **I/II/III (%)** | 13.51/35.14/51.35 | 8.82/44.12/47.06 | - |

Fa, CRC patients before treatment; Fb, CRC patients after surgery.

We obtained a total of 4 992 311 16S rDNA sequences from 107 stool samples, with an average of 46 657 ± 2955 reads per sample in the whole cohort.

We generated 1108 total operational taxonomic units (OTUs) at a 97 % similarity level, with an average of 244 ± 67, 190 ± 74, 130 ± 60 OTUs in the Normal, Fa, and Fb groups, respectively. The maximum number of OTUs for a single sample was 401, whereas the minimum, which was found in the Fb group, was only 27 (**Table S1**). There were 495 common OTUs in all groups, with the Fa group having the most specific OTUs, whereas the Fb group having the least specific OTUs (**Figure S1**).

### Richness and diversity

The observed species and Chao richness index, Shannon, and Simpson diversity index were used to describe the alpha diversity features of the bacterial communities in our samples. We observed that the species richness and diversity in Fa and Fb were decreased compared with those in the Normal group. A strong decrease in biodiversity was observed in the Fb group, especially in the stages II and III subgroup of Fb, compared with that in the other groups (**Figures 1A, S2**).

**Figure 1.**
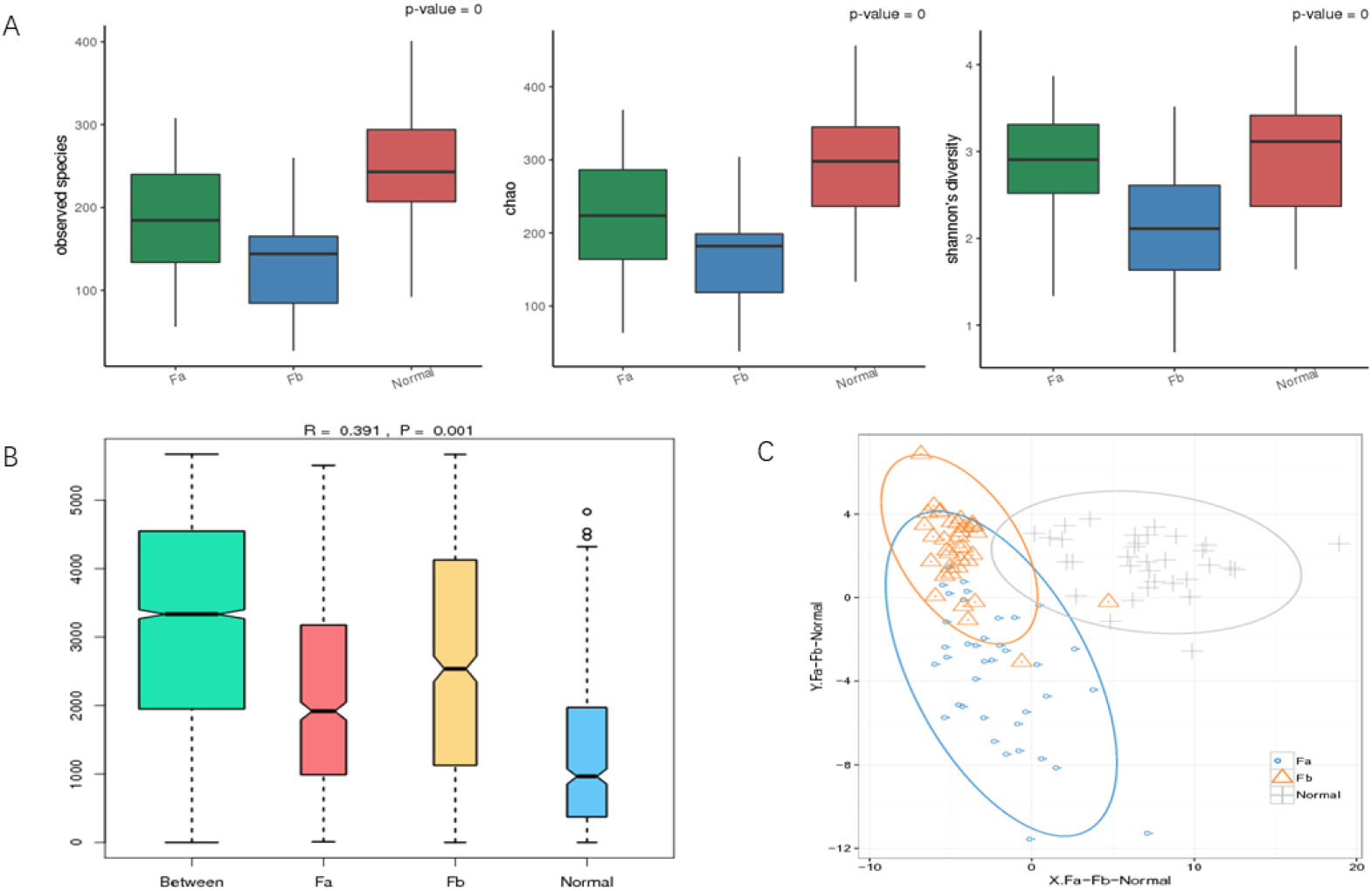
Microbiota biodiversity. (A) Alpha diversity. Bigger indexes of the observed species and Chao reflect greater richness, whereas a bigger Shannon index reflects greater diversity. (B) Similarities. “Between” shows the differences between groups, whereas the rest show differences within groups, R > 0 indicates differences between groups were more obvious than within groups, P value < 0.01 indicates significance. (C) OTUs based PLSDA. ADONIS, Fa-Normal, R^2^ = 0.11797, P = 0.001 (***); Fa-Fb, R^2^ = 0.05057, P = 0.001(***); Fb-Normal, R^2^ = 0.18593, P = 0.001(***). OTU, operational taxonomic unit; PLSDA, partial least squares discrimination analysis; Fa, CRC patients before treatment; Fb, CRC patients after surgery.

We used analysis of similarities (ANOSIM) to estimate the similarity among groups. Our results indicated that differences among groups were more significant than differences within groups (R-value = 0.164, P = 0.001) (**Figure 1B**). At the same time, a beta diversity evaluation, represented by partial least squares discrimination analysis (PLSDA), showed a clustering of samples according to disease with an obvious separation between the Fa and Normal groups, and the Fb and Normal groups, but a partial separation between the Fa and Fb groups. Permutational multivariate analysis of variance confirmed this observation (ADONIS, Fa-Normal, R^2^ = 0.11797, P = 0.001; Fa-Fb, R^2^ = 0.05057, P = 0.001; Fb-Normal, R^2^ = 0.18593, P = 0.001) (**Figure 1C**).

### Bacterial microbiota composition

We analyzed the composition and abundance of bacteria at all taxonomic levels. As expected, we found that a large majority of the bacteria in the Fa, Fb, and normal samples belonged to the phyla Bacteroidetes, Firmicutes, Proteobacteria, Fusobacteria, and Actinobacteria. We further identified that the distribution of the major phyla in the Normal group was consistent with published data. Further comparison of the relative abundance revealed clear differences. The most abundant phylum in the Normal group was Bacteroidetes, followed by Firmicutes, Proteobacteria, and Actinobacteria. However, an increase in the distribution of Proteobacteria, Fusobacteria, and Verrucomicrobia was observed in the Fa group. Except for an increase in the distribution of Fusobacteria and Verrucomicrobia, the Fb group was characterized by a notable increase in Proteobacteria and a decrease in Firmicutes (**Figure S3**).

We identified a total of 173 genera at the genus level, with the dominant genus among all groups being *Bacteroides* (34.89 %, 32.34 %, and 22.89 % in Normal, Fa, and Fb groups, respectively). However, apart from this, the composition and prevalence of genera was different among the three groups. In the Normal group, we identified the following genera: *Prevotella* (21.49 %), *Faecalibacterium* (8.58 %), *Roseburia* (3.28 %), and *Ruminococcus* (3.02 %). The Fa group was characterized by the presence of *Escherichia* (10.69 %), *Faecalibacterium* (5.49 %), *Prevotella* (4.78 %)*, and Parabacteroides* (3.84 %); whereas in the Fb group, we observed *Escherichia* (18.56 %), *Parabacteroides* (6.81 %), *Enterococcus* (5.82 %)*, and Morganella* (4.68 %) (**Table S2**). Accordingly, *Escherichia* belonging to the phylum Proteobacteria, *Fusobacterium* belonging to Fusobacteria, and *Parabacteroides* were found to be enriched in CRC patients (Fa and Fb groups) compared with that in the Normal group, whereas *Prevotella* was demonstrated to be overrepresented in the Normal group. The presence of *Faecalibacterium* was scarce, whereas *Enterococcus* was abundant in the Fb group (**Figure 2**). Although *Bacteroides* exhibited a similar relative abundance at the genus level, at the species level, the abundance of *B. fragilis* was shown to vary among groups (**Figure S4**), being enriched in the Fa and Fb groups.

**Figure 2.**
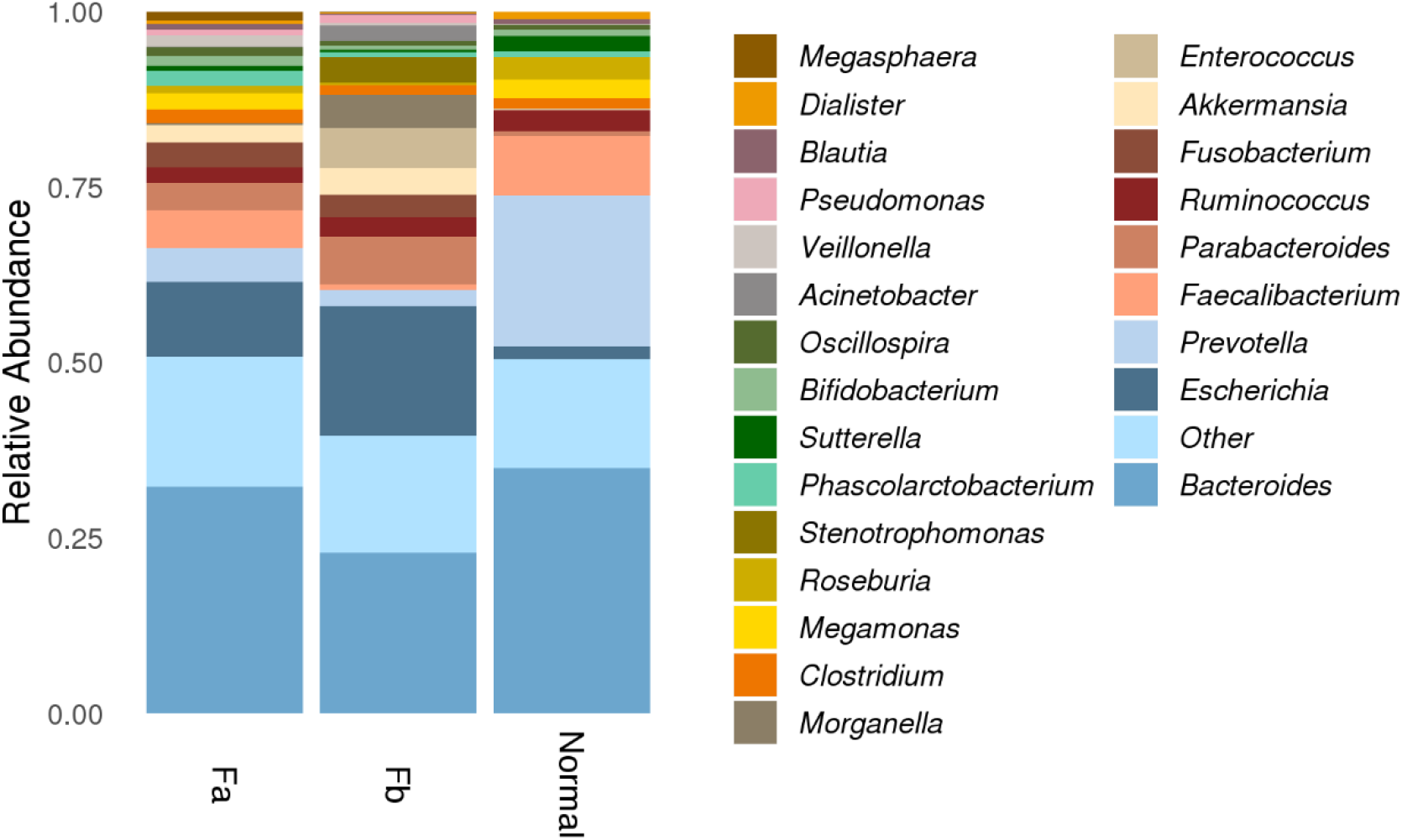
Microbiota composition in every group at the genus level. Relative abundances of less than 0.5 % were combined and shown as other. Fa, CRC patients before treatment; Fb, CRC patients after surgery.

### Identification of differential microbes and key taxa (biomarkers)

To identify the key bacteria causing divergence between different groups, we used the linear discriminant analysis (LDA) effect size (LEfSe) biomarker discovery tool, which could compare two or more groups, and search for biomarkers showing statistical differences. We performed LEfSe analysis at both the family and genus levels, and found 52 discriminative features using a threshold of LDA score of 2 (P value <0.01) at the genus level (**Table S3**). We observed that *Prevotella* (LDA = 4.95, P < 0.01), *Faecalibacterium* (LDA = 4.66, P < 0.001), *Roseburia* (LDA = 4.23, P < 0.001), *Megamonas* (LDA = 4.21, P < 0.001), and *Sutterella* (LDA = 4.08, P < 0.001) were the dominant microbes in the Normal group; *Fusobacterium* (LDA = 4.22, P < 0.001) was the dominant genus in the Fa group, whereas *Escherichia* (LDA = 4.96, P < 0.001), *Enterococcus* (LDA = 4.49, P < 0.001), and *Stenotrophomonas* (LDA = 4.17, P < 0.01) were the dominant genera in the Fb group. The genera with an LDA score higher than 3 are displayed in **Figure 3A**. To further compare the relative abundance of these primary biomarkers in all groups, we evaluated the average relative abundance of bacteria with an LDA score higher than 4 in every group. Except for *Bacteroides*, all other bacteria showed significant differences (P < 0.01) (**Figure 3B**).

**Figure 3.**
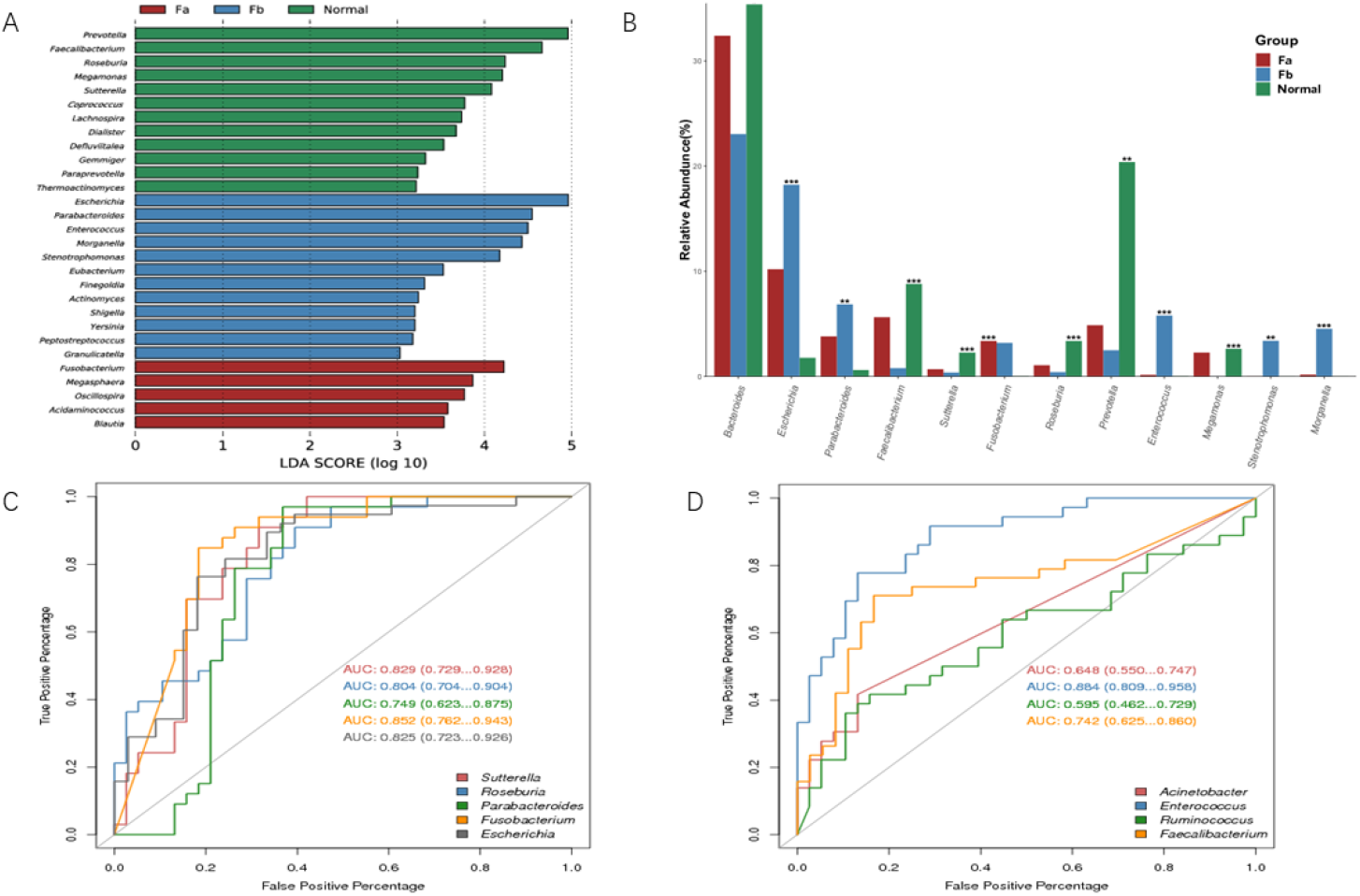
A) LEfSe analysis for taxonomic biomarkers on the genus level among the three groups. Each color represents one group. (B) Differential comparison of key microbes. ***, P < 0.001; **, 0.001 ≤ P ≤ 0.01; *, 0.01 ≤ P ≤ 0.05. (C) ROC curves for evaluation of the classification model between Fa and normal samples. (D) ROC curves for evaluation of the classification model between Fa and Fb samples. LEfSe, linear discriminant analysis effect size; ROC, receiver operating characteristic; AUC, area under the ROC curve; Fa, CRC patients before treatment; Fb, CRC patients after surgery.

Consecutively, to explore whether these differential microbes were suitable for CRC detection, or classification of CRC samples before or after treatment, we used the receiver operating characteristic (ROC) curve to evaluate their predictive power. First, we calculated the area under the ROC curve (AUC) of the microbes between the Fa and Normal groups, and found that the most discriminative genus was that of *Fusobacterium* with an AUC of 0.852 (**Figure 3C**). Consistent with previous studies suggesting *Fusobacterium* as prevalent in the gut of CRC patients^11,12^, potentially accelerating tumorigenesis^10^, our results further confirmed this enrichment and indicated a potential biomarker for detecting CRC. Following, we evaluated the classification model comparing samples from before (Fa group) and after (Fb group) treatment. Our results revealed that the most discriminative microbe, which was shown to be enriched in the Fb group, with an AUC value of 0.884, was *Enterococcus* **(Figure 3D**).

### Function analysis and correlation with clinic data

We used Picrust2 to predict the MetaCyc pathways of microbiota in every sample. This analysis revealed the differential functions between CRC patients and healthy individuals and between CRC patients before and after treatment. We observed that the pathways enriched in the Fa group compared with those of the Normal group were those of lipid, fatty acid, amino acid, aldehyde, alcohol, and aromatic compound degradation **(Figure 4A**). Only the metabolic regulator biosynthesis as well as fatty acid and lipid degradation pathways were identified to be significantly (P < 0.01, |log2FC| > 1) different in the Fa and Fb groups. (**Figure 4B**).

**Figure 4.**
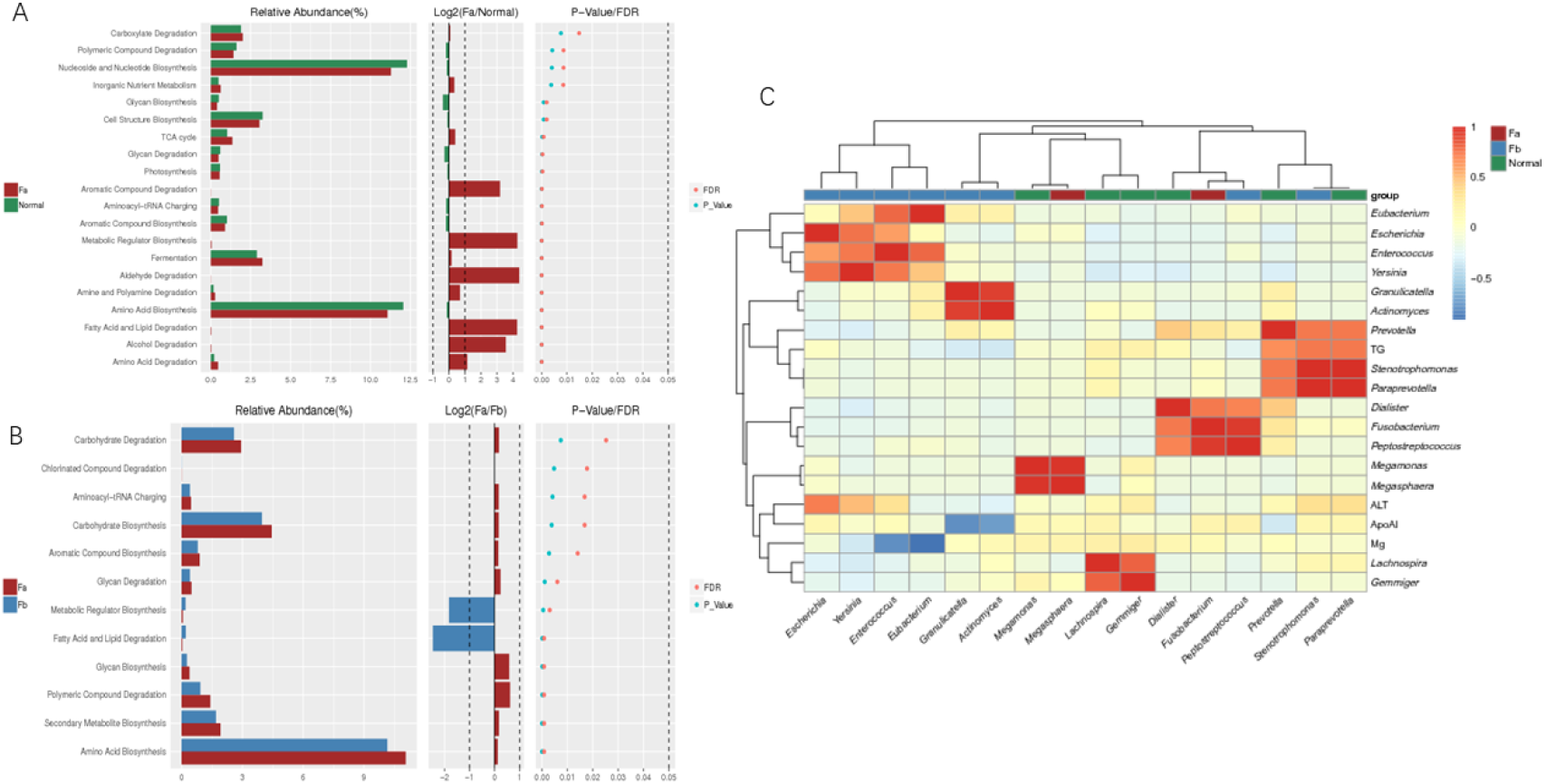
(A) Comparison of differential functional pathways in the Fa and Normal groups. (B) Comparison of differential functional pathways in the Fa and Fb groups. (C) Heatmap of correlation. Fa, CRC patients before treatment; Fb, CRC patients after surgery.

We collected 56 clinical indexes of CRC patients, including biochemical criteria and routine blood examinations. At the genus level, 27 differentially abundant microbes (LDA score > 3, P < 0.01) were selected and related to the clinical data. We calculated the Pearson coefficient of pairwise correlation between microbes and clinical indexes, and characters exhibiting a high correlation (Pearson coefficient ≥ 0.7) are displayed on a heatmap **(Figure 4C**). A similar abundance model and very strong correlation could be observed in some microbes, such as *Escherichia, Enterococcus, Yersinia*, and *Eubacterium*, which were enriched and clustered well in the Fb group, whereas *Fusobacterim* and *Peptostreptococcus* were found to be clustered together, with a Pearson coefficient of 0.95. We selected ALT, ApoA1, Mg^2+^, and TG, among all the clinical indexes. ALT was shown to be positively related to *Escherichia* (R = 0.7303), TG was positively related to *Stenotrophomonas* (R = 0.7333) and *Paraprevotella* (R = 0.7328), whereas Mg^2+^ was negatively related to *Eubacterium* (R = 0.9096) and *Enterococcus* (R = 0.7795), and ApoA1 was negatively related to *Granulicatella* (R = 0.7785).

### Biomarkers and prognosis

We followed-up 32 CRC patients who had provided pre- and post-treatment stool samples, and recorded their current living state (approximately 3 y after surgery). We evaluated the changes in important bacteria in paired stool samples. We observed a decrease in the presence of *Fusobacterium* in most patients treated at the SUN YAT-SEN University Cancer Center after surgery, except for two cases of distinct increases. One CRC patient died after surgery, while the other had chronic enteritis (**Figure 5A**). Meanwhile, most patients treated at the SUN YAT-SEN Memorial Hospital exhibited the same decrease in *Fusobacterium* after surgery (**Figure 5B**). Five patients (A147, A119, B103, B106, B112) either developed postoperative recidivation or died. We observed that most of the samples showed an abnormal increase in *Fusobacterium*, and all of them exhibited an obvious decrease in beneficial bacteria (*Faecalibacterium* and *Prevotella*) (**Figure 5**). This finding suggested that an abnormal increase in *Fusobacterium* and a distinct reduction of probiotic might indicate poor prognosis. This supposition should be followed by a large cohort and more stages of postoperative sampling, such as one month after surgery, three months after surgery, and so on.

**Figure 5.**
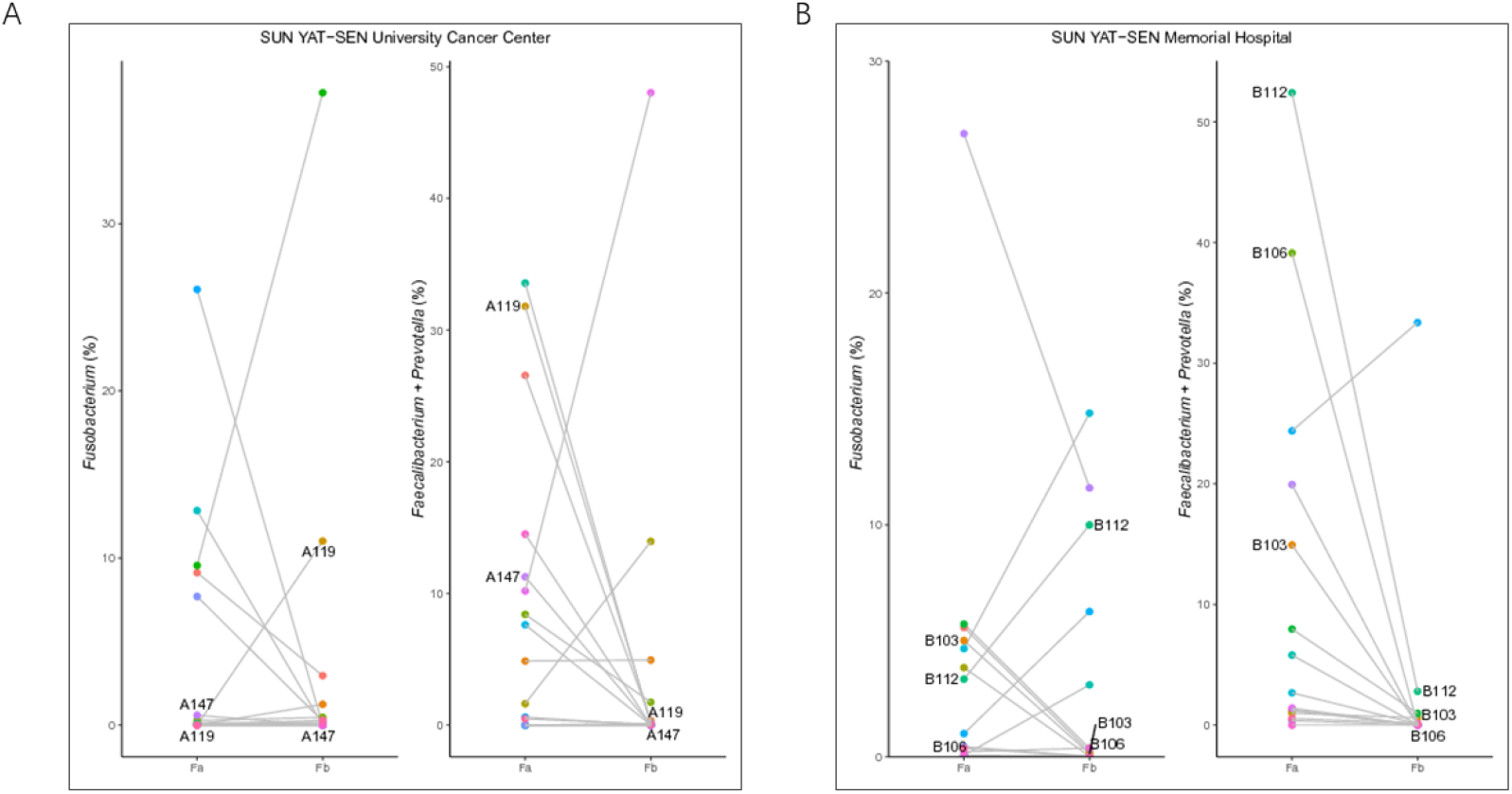
Relative abundance of *Fusobacterium, Faecalibacterium*, and *Prevotella* in every paired sample before and after surgery. (A) Samples collected from the SUN YAT-SEN University Cancer Center. (B) Samples collected from the SUN YAT-SEN Memorial Hospital. Fa, CRC patients before treatment; Fb, CRC patients after surgery.

## 3. Discussion

In this study, we compared the fecal microbiota of CRC patients to those of healthy individuals and CRC patients that underwent surgery. We observed changes in the microbiota in all three groups. Moreover, the richness and biodiversity among these groups was found to differ. In particular, we found a decrease in biodiversity in the Fa group, and a strong decrease in biodiversity in CRC patients who had undergone surgical operation (Fb group, approximately 1 wk after surgery), indicating that surgery might lead to serious microbiota dysbiosis. However, as patients in the Fb group were administered antibiotics after surgery, we could not eliminate the effect of antibiotics on biodiversity.

We also observed obvious differences in bacterial composition. The composition of fecal microbiota in the Fa and Fb groups was clearly different from the microbiota of healthy individuals, with a clear increase in the abundance of *Escherichia, Parabacteroides*, and *Fusobacterium* being observed in both the Fa and Fb groups. Previous studies have reported a number of microbial species found in CRC patients, most of which were present in our dataset as well. For instance, *Fusobacterium* was reported to coexist with tumors, and was considered to positively regulate tumor cell propagation^12,13^. *F. nucleatum* was demonstrated to increase the tumor burden and selectively expand myeloid derived immune cells, such as CD11b^+^, and myeloid derived suppressor cells in an Apc^Min/+^ mouse model^10^. Other studies suggested that through the recruitment of tumor-infiltrating immune cells, *Fusobacteria* might generate a proinflammatory microenvironment that is conducive for colorectal neoplasia progression. In accordance with these findings, our results showed that *Fusobacterium* was identified to be the principal genus in the Fa group. Moreover, the classification model between the Fa and Normal groups was credible with an AUC of 0.852, suggesting that *Fusobacterium* could be used as a potential biomarker for CRC patients. Thus, the increased abundance of *Fusobacterium* could be linked with a high risk of CRC. Enterotoxigenic *B. fragilis* has been identified as a potential driver of CRC in both human and mouse studies^18,19,20^. The toxin of *B. fragilis* is known to cause human inflammatory diarrhea. However, it can also asymptomatically colonize a proportion of the human population, thereby triggering colitis and strongly inducing colonic tumors via activation of the T-helper type 17 T-cell responses^20^. In our dataset, we found that the species of *B. fragilis* was prominent in CRC patients either before or after surgery, especially in the Fb group*. E. coli* is a commensal bacterium of the human gut microbiota, but some pathogenic strains have acquired the ability to induce chronic inflammation or produce toxins, such as cyclomodulins, which could participate in carcinogenesis processes^1,21,22^. We also tested the enrichment of *Escherichia* in CRC patients, especially in the Fb group. At the species level, the presence of *E. coli* was shown to be obviously increased in the Fa and Fb groups than in the Normal group. All the aforementioned bacteria involved in CRC are known to be proinflammatory-associated, and hence might colonize faster in an inflammatory conducive environment. As the fecal samples of the Fb group in this study were obtained from CRC patients who had recently undergone surgery, their intestinal microenvironments were probably unstable, with some potentially exhibiting an inflammatory response, and some may have bad prognosis. Therefore, our findings that the abundance of *B. fragilis* and *E. coli* was the largest in the Fb group, whereas *Fusobacterium* showed a slight decrease compared with that in the Fa group, was justified. The distribution trend of these bacteria after surgery should be analyzed in different stages post-surgery, and thus further research is needed.

*Fa. prausnitzii* is one of the most abundant bacteria in the human intestinal microbiota of healthy individuals, and the most important butyrate-producing bacteria in the human colon^23^, representing more than 5 % of the total bacterial population^16^. Further, this bacterium has shown potential to function as a probiotic in the treatment of Crohn’s disease^24^. Changes in the abundance of *Fa. prausnitzii* have been linked to dysbiosis in several human disorders. To date, this commensal bacterium has been considered as a bioindicator of human health. *Prevotella*, a commensal bacterial genus known to produce short chain fatty acids and that possesses potent anti-inflammatory effects, has been reported to be more commonly found in non-Westerners, who prefer a plant-rich diet^25,26^. Studies have confirmed that maternal carriage of *Prevotella* during pregnancy was associated with protection against food allergies in the offspring^26^. In our study, the numbers of *Faecalibacterium* and *Prevotella* were reduced in CRC patients, especially in the Fb group. We further noted that this reduction was more notable in patients who recrudesced or died after surgery.

The genus *Enterococcus* is of great relevance to human health because of its role as a major causative agent of healthcare-associated infections; it includes resilient and versatile species able to survive under harsh conditions, most demonstrating intrinsic resistance to common antibiotics, such as virtually all cephalosporins, aminoglycosides and clindamycin^27^. As individuals in the Fb group had recently undergone resection, and were administered antibiotics, serious microbial dysbiosis might have occurred in their gut. As *Enterococcus* is known to exhibit versatility and drug-resistance, it could still adapt to the post-operation environment. Thus, the increase in the numbers of this bacteria in the Fb group was justified.

## 4. Conclusions

In conclusion, this study showed the different composition of fecal microbiota among Chinese healthy individuals and CRC patients before and after surgery. We also identified *Fusobacterium* as a potential biomarker for CRC screening. We also found changes in the numbers of *Fusobacterium*, *Faecalibacterium*, and *Prevotella*, which were shown to be related to prognosis after surgery. This finding could contribute to CRC early screening and prognosis monitoring in Chinese or Asian populations, in combination with multiple factors of cfDNA methylation and alteration.

## 5. Materials and Methods

### Samples

CRC patients were selected from Sun Yat-sen University Cancer Center and Sun Yat-sen Memorial Hospital, China. All CRC patients were diagnosed according to endoscopic and histological parameters. None of the patients had undergone any treatment, such as radiotherapy or chemotherapy, before enrollment. Exclusion criteria included other neoplasms, other tumor history, tuberculosis, infection by hepatitis B (HBV), hepatitis C (HCV), or HIV, and the use of antibiotics 1 mo prior to hospitalization. The cancer stage was identified according to the TNM classification of malignant tumors. CRC patients were first enrolled and hospitalized; then, their fecal samples were collected before surgery (approximately 2-3 d before surgery). CRC patients underwent operative treatment, then administered cefminox sodium for 3 d in a row. Following collection of their fecal samples approximately 1 wk after resection surgery, they were allowed to leave the hospital. Normal samples were collected from healthy individuals with no history of cancer, chronic enteritis, chronic constipation, bloody stools, chronic appendicitis, or chronic cholecystitis.

In total, 107 stool samples were collected and divided into the following three groups: 33 samples from healthy individuals, named Normal group; 38 samples from CRC patients before surgery, named Fa group; 36 samples from CRC patients approximately 1 wk after surgery, named Fb group. Basic information regarding this study population is listed in **Table 1**. Among them, 32-paired samples, each paired with pre- and post-treatment stool samples was obtained from the same patient. The current survival state of these 32 patients was recorded (**Table S4**).

Clinical data were also collected simultaneously. In total, 56 clinical indexes of CRC patients (pre- and post-surgery corresponding to 72 stool samples) were collected, including biochemical criteria and routine blood examinations.

### Sample preparation and genomic DNA extraction

None of the patients were subjected to any invasive operation, such as endoscopic or clyster, at least 5 d before sampling. A light diet was suggested 3 d before sampling. About 5 g of fresh stool samples were collected by patients themselves immediately after defecation using stool collection devices, and then shipped on dry ice in insulated containers to a central lab, where the samples were immediately stored at −80 °C until further processing. The microbiota DNA was extracted as previously described^28^. Samples were treated with lysozyme, proteinase K, and SDS, then purified with phenol-chloroform-isoamylalcohol, precipitated using glycogen, sodium acetate and cold isopropanol, washed with 75% ethanol and resuspended in 1× TE buffer. DNA integrity and purification were detected by agarose gel electrophoresis (1 %, 150 V, 40 min).

### Library construction and next generation sequencing (NGS)

Qualified samples were used for the library preparation process. The microbiota DNA was amplified by polymerase chain reaction (PCR) with a bacterial 16S rDNA V4 region universe primer pair (515F: 5’-GTGCCAGCMGCCGCGGTAA-3’ and 806R: 5’-GGACTACHVGGGTWTCTAAT-3’). PCR was performed using the following conditions: 3 min denaturation at 94 °C; 25 cycles of denaturation at 94 °C for 45 s, annealing at 50 °C for 60 s, elongation at 72 °C for 90 s; and final extension at 72 °C for 10 min. The PCR products were purified using AMPure XP beads (Axygen). Barcoded libraries were generated by emulsion PCR and quantitated in the following two ways: the average molecule length was determined using the Agilent 2100 bioanalyzer instrument and the library was quantified by real-time quantitative PCR (QPCR).

The qualified libraries were sequenced using the Illumina HiSeq2500 platform with the PE250 sequencing strategy (PE251 + 8 + 8 + 251).

### Sequence processing

Raw sequences were assigned to each sample based on their unique barcode and primer; subsequently, the barcodes and primers were removed. At the same time, paired-end low-quality reads were filtered based on quality score, adapter contamination, and N base ratio.

Paired-end clean reads were merged using FLASH (fast length adjustment of short reads, v1.2.11)^29^ according to the relationship of the overlap between paired-end reads. This was done when at least 15 bp of the read overlapped the read generated from the opposite end of the same DNA fragment, the maximum allowable error ratio of an overlap region was set as 0.1, and merged sequences were called clean tags.

Tags were assigned to OTUs using USEARCH (v7.0.1090) software^30^, and tags with ≥ 97 % similarity were clustered to the same OTU. It has been reported that a singleton OTU could be obtained due to sequencing errors or chimeras generated during PCR; therefore, chimeric sequences were detected and removed using UCHIME (v4.2.40)^31^ according to the match of representative OTUs to the gold database (v20110519). The abundance of each OTU was quantified using usearch_global algorithm by matching all clean tags to final OTUs, and normalized using a standard number corresponding to the sample with the least sequences.

Representative OTUs were annotated using the RDP classifier (v2.2) software^32^ based on the homolog of the Greengene database (v201305), with the confidence threshold set to 0.8. OTUs without annotation or annotated to polluted species were removed, and the number of effective tags and information regarding OTU taxonomic synthesis were recorded in a table for the next analysis. The structure of the bacterial community of each sample was analyzed at all levels of taxonomy, with the relative abundance less than 0.5 % in all samples combined with others.

### Statistical Analysis

Common and specific OTUs among groups were compared and displayed using VennDiagram R (v3.1.1). Analysis of similarities were performed using Bray-Curtis in the vegan package of R (v3.5.1); comparison of differences between and within the groups was available, thereby allowing testing of the availability of grouping.

Alpha diversity was applied to analyze the complexity of species diversity of a sample using many indexes, such as observed species, Chao, Ace, Shannon, and Simpson. All indices of our samples were calculated using Mother (v1.31.2)^33^, and comparisons among groups were performed using the Kruskal test. Observed species and Chao were selected to identify community richness, whereas Shannon was used to identify community diversity. Beta diversity^34,35^ was used to evaluate the differences in species complexity among different samples, and was calculated on both weighted and unweighted UniFrac using QIIME (v1.80). Partial least squares discrimination analysis (PLS-DA) was built using the mixOmics library of R (v3.2.1), which was used to estimate the classification of samples and assess the variation in study groups.

We analyzed the differential abundance at the phylum, class, order, family, genus, and species levels. Differential abundance analysis was performed using LEfSe^36^, with the P value less than 0.01 and an LDA score more than 2 being considered significant. To quantify the effective size of the differential taxa, we used the fold change of the mean relative abundance between groups. Comparisons between probabilities, as well as overall differences in the mean relative abundance of each taxon between the two groups were evaluated using a paired Wilcoxon rank sum test. Comparisons among three or more groups were performed using the Kruskal-Wallis test.

The ROC curve was used to assess the confidence level of the classification model. Accordingly, ROC analysis and the AUC values were calculated using the pROC package of R.

MetaCyc pathway prediction was performed using Picrust2, MetaCyc (https://metacyc.org/) containing pathways involved in primary and secondary metabolism, related metabolites, and enzymatic reactions. Differential functions were analyzed using the Wilcox-test between the two groups. Correlation was tested by Pearson’s coefficient using the R package.

## Abbreviations

CRC: colorectal cancer
ROS: reactive oxidative species
OTUs: operational taxonomic units
LDA: linear discriminant analysis
LEfSe: linear discriminant analysis effect size
ROC: receiver operating characteristic
AUC: area under the ROC curve
NGS: next generation sequencing
PCR: polymerase chain reaction
QPCR: quantitative PCR
FLASH: fast length adjustment of short reads
PLS-DA: partial least squares discrimination analysis
ALT: alanine transaminase
ApoA1: apolipoprotein A1
TG: triglyceride

## Acknowledgements

We acknowledge the volunteers who participated in our study.

## Authors’ contributions

Qiulin Yao: Methodology, Data curation, Formal analysis, Visualization, Writing-Original draft preparation. Meifang Tang: Conceptualization, Methodology, Supervision, Writing-Review & Editing. Liuhong Zeng: Methodology, Selecting samples, Communication. Zhonghua Chu: Conceptualization, Resources. Hui Sheng: Resources. Yuyu Zhang: Resources. Yuan Zhou: Library preparation. Hongyun Zhang: Review. Huayan Jiang: Providing information. Mingzhi Ye: Funding acquisition, Resources.

## Availability of data and materials

The data reported in this study are also available in the CNGB Nucleotide Sequence Archive (CNSA: https://db.cngb.org/cnsa; accession number CNP0001385).

## Ethics approval and consent to participate

The study was approved by appropriate Institutional Review Boards (IRB) of the BGI (NO. BGI-IRB15100-T1).

## Consent for publication

Not applicable.

## Author details

^1^Clinical laboratory of BGI Health, BGI-Shenzhen, Shenzhen 518083, China. ^2^BGI Education Center, University of Chinese Academy of Sciences, Shenzhen 518083, China. ^3^Guangdong Provincial Key Laboratory of Malignant Tumor Epigenetics and Gene Regulation, Department of Gastrointestinal Surgery, Sun Yat-sen Memorial Hospital, Sun Yat-sen University, Guangzhou 510060, China. ^4^Department of Experimental Research, State Key Laboratory of Oncology in South China, Collaborative Innovation Center for Cancer Medicine, Sun Yat-Sen University Cancer Center, Guangzhou 510060, China. ^5^BGI Genomics, BGI-Shenzhen, Shenzhen 518083, China. ^6^BGI-Guangzhou Medical Laboratory, BGI-Shenzhen, Guangzhou 510006, China

