## Supplementary for "Human fecal microbiota is associated with colorectal cancer"

**Table S1 OTUs stat of all groups**

|  | Fa | Fb | Normal |
| --- | --- | --- | --- |
| Sample n | 38 | 36 | 33 |
| Min OTU n | 56 | 27 | 92 |
| Max OTU n | 308 | 260 | 401 |
| Mean OTU (±SD） | 190.5±73.7 | 130.7±60.0 | 244.5±67.1 |

**Table S2 Bacterial microbiota composition of each group at the genus level**

| Taxon | Fa | Fb | Normal |
| --- | --- | --- | --- |
| Bacteroides | 0.3233553 | 0.228916 | 0.3489311 |
| Other | 0.1845805 | 0.166014 | 0.1560167 |
| Escherichia | 0.1069364 | 0.185643 | 0.017645 |
| Faecalibacterium | 0.0549211 | 0.007359 | 0.0858073 |
| Prevotella | 0.0477655 | 0.023284 | 0.2148762 |
| Parabacteroides | 0.0383737 | 0.068063 | 0.0059306 |
| Fusobacterium | 0.0348777 | 0.032025 | 0.0003008 |
| Akkermansia | 0.0245388 | 0.037431 | 0.0014791 |
| Phascolarctobacterium | 0.0224098 | 0.006459 | 0.0073193 |
| Megamonas | 0.0222687 | 5.12E-05 | 0.0264367 |
| Ruminococcus | 0.0221821 | 0.027505 | 0.0301713 |
| Clostridium | 0.0205138 | 0.01398 | 0.0148241 |
| Veillonella | 0.0165846 | 0.004129 | 0.0007382 |
| Bifidobacterium | 0.0142193 | 0.005317 | 0.0094924 |
| Oscillospira | 0.0127759 | 0.00792 | 0.0063459 |
| Megasphaera | 0.0125637 | 0.001645 | 0.0005473 |
| Roseburia | 0.0100033 | 0.003841 | 0.0328248 |
| Pseudomonas | 0.0083752 | 0.011202 | 0.0000355 |
| Blautia | 0.0078228 | 0.001616 | 0.0071123 |
| Sutterella | 0.0067414 | 0.00353 | 0.0224923 |
| Dialister | 0.004987 | 0.001541 | 0.0102567 |
| Morganella | 0.0016505 | 0.046792 | 0.0000134 |
| Enterococcus | 0.001483 | 0.058167 | 0.0003229 |
| Acinetobacter | 6.827E-05 | 0.021133 | 7.838E-05 |
| Stenotrophomonas | 1.72E-06 | 0.036436 | 2.01E-06 |

**Table S3 Feature list of LefSe analysis**

| Biomarker | Logarithm value | groups | LDA score | P-value |
| --- | --- | --- | --- | --- |
| Bacteria.Fusobacteria.Fusobacteriia.Fusobacteriales.Fusobacteriaceae.Fusobacterium | 4.5642 | Fa | 4.2233 | 2.71E-07 |
| Bacteria.Firmicutes.Clostridia.Clostridiales.Veillonellaceae.Megasphaera | 4.1209 | Fa | 3.8634 | 0.006373 |
| Bacteria.Firmicutes.Clostridia.Clostridiales.Ruminococcaceae.Oscillospira | 4.2452 | Fa | 3.7680 | 0.009607 |
| Bacteria.Firmicutes.Clostridia.Clostridiales.Veillonellaceae.Acidaminococcus | 3.8544 | Fa | 3.5794 | 0.002237 |
| Bacteria.Firmicutes.Clostridia.Clostridiales.Lachnospiraceae.Blautia | 3.9529 | Fa | 3.5340 | 3.23E-07 |
| Bacteria.Firmicutes.Clostridia.Clostridiales.Tissierellaceae.Parvimonas | 3.1618 | Fa | 2.8919 | 2.61E-06 |
| Bacteria.Actinobacteria.Coriobacteriia.Coriobacteriales.Coriobacteriaceae.Collinsella | 3.4166 | Fa | 2.8680 | 0.007137 |
| Bacteria.Firmicutes.Erysipelotrichi.Erysipelotrichales.Erysipelotrichaceae.Bulleidia | 2.4685 | Fa | 2.3687 | 0.00015 |
| Bacteria.Firmicutes.Clostridia.Clostridiales.Lachnospiraceae.Lachnobacterium | 2.5433 | Fa | 2.3459 | 3.46E-08 |
| Bacteria.Proteobacteria.Gammaproteobacteria.Enterobacteriales.Enterobacteriaceae.Escherichia | 5.3149 | Fb | 4.9589 | 2.21E-06 |
| Bacteria.Bacteroidetes.Bacteroidia.Bacteroidales.Porphyromonadaceae.Parabacteroides | 4.8723 | Fb | 4.5449 | 0.002415 |
| Bacteria.Firmicutes.Bacilli.Lactobacillales.Enterococcaceae.Enterococcus | 4.8049 | Fb | 4.4965 | 2.69E-12 |
| Bacteria.Proteobacteria.Gammaproteobacteria.Enterobacteriales.Enterobacteriaceae.Morganella | 4.6980 | Fb | 4.4286 | 4.76E-05 |
| Bacteria.Proteobacteria.Gammaproteobacteria.Xanthomonadales.Xanthomonadaceae.Stenotrophomonas | 4.5283 | Fb | 4.1715 | 0.007819 |
| Bacteria.Firmicutes.Erysipelotrichi.Erysipelotrichales.Erysipelotrichaceae.Eubacterium | 3.8985 | Fb | 3.5270 | 1.04E-07 |
| Bacteria.Firmicutes.Clostridia.Clostridiales.Tissierellaceae.Finegoldia | 1.8628 | Fb | 3.3112 | 0.000734 |
| Bacteria.Actinobacteria.Actinobacteria.Actinomycetales.Actinomycetaceae.Actinomyces | 3.5457 | Fb | 3.2399 | 2.52E-07 |
| Bacteria.Proteobacteria.Gammaproteobacteria.Enterobacteriales.Enterobacteriaceae.Shigella | 1.4266 | Fb | 3.2021 | 0.00013 |
| Bacteria.Proteobacteria.Gammaproteobacteria.Enterobacteriales.Enterobacteriaceae.Yersinia | 3.5899 | Fb | 3.2015 | 0.005547 |
| Bacteria.Firmicutes.Clostridia.Clostridiales.Peptostreptococcaceae.Peptostreptococcus | 3.4937 | Fb | 3.1772 | 1.59E-08 |
| Bacteria.Firmicutes.Bacilli.Lactobacillales.Carnobacteriaceae.Granulicatella | 3.3126 | Fb | 3.0328 | 9.06E-07 |
| Bacteria.Actinobacteria.Coriobacteriia.Coriobacteriales.Coriobacteriaceae.Atopobium | 3.1918 | Fb | 2.8886 | 0.001358 |
| Bacteria.Actinobacteria.Actinobacteria.Actinomycetales.Corynebacteriaceae.Corynebacterium | 2.1291 | Fb | 2.7732 | 0.001681 |
| Bacteria.Actinobacteria.Coriobacteriia.Coriobacteriales.Coriobacteriaceae.Eggerthella | 3.0758 | Fb | 2.7113 | 5.81E-06 |
| Bacteria.Actinobacteria.Actinobacteria.Actinomycetales.Micrococcaceae.Rothia | 2.9697 | Fb | 2.6539 | 0.009373 |
| Bacteria.Firmicutes.Bacilli.Lactobacillales.Aerococcaceae.Abiotrophia | 2.8618 | Fb | 2.5707 | 0.003524 |
| Bacteria.Firmicutes.Clostridia.Clostridiales.Eubacteriaceae.Pseudoramibacter_Eubacterium | 2.9035 | Fb | 2.5650 | 1.52E-05 |
| Bacteria.Proteobacteria.Betaproteobacteria.Neisseriales.Neisseriaceae.Eikenella | 2.8374 | Fb | 2.5419 | 0.002865 |
| Bacteria.Firmicutes.Clostridia.Clostridiales.Mogibacteriaceae.Mogibacterium | 1.9769 | Fb | 2.3139 | 8.53E-05 |
| Bacteria.Bacteroidetes.Bacteroidia.Bacteroidales.Prevotellaceae.Prevotella | 5.3543 | Normal | 4.9549 | 0.001556 |
| Bacteria.Firmicutes.Clostridia.Clostridiales.Ruminococcaceae.Faecalibacterium | 5.0110 | Normal | 4.6587 | 3.91E-10 |
| Bacteria.Firmicutes.Clostridia.Clostridiales.Lachnospiraceae.Roseburia | 4.5880 | Normal | 4.2357 | 7.99E-11 |
| Bacteria.Firmicutes.Clostridia.Clostridiales.Veillonellaceae.Megamonas | 4.4381 | Normal | 4.2059 | 3.47E-06 |
| Bacteria.Proteobacteria.Betaproteobacteria.Burkholderiales.Alcaligenaceae.Sutterella | 4.4054 | Normal | 4.0795 | 2.55E-09 |
| Bacteria.Firmicutes.Clostridia.Clostridiales.Lachnospiraceae.Coprococcus | 4.1063 | Normal | 3.7730 | 8.55E-12 |
| Bacteria.Firmicutes.Clostridia.Clostridiales.Lachnospiraceae.Lachnospira | 4.0642 | Normal | 3.7363 | 1.02E-13 |
| Bacteria.Firmicutes.Clostridia.Clostridiales.Veillonellaceae.Dialister | 4.0510 | Normal | 3.6748 | 7.56E-05 |
| Bacteria.Firmicutes.Clostridia.Clostridiales.Lachnospiraceae.Defluviitalea | 1.3330 | Normal | 3.5328 | 0.009918 |
| Bacteria.Firmicutes.Clostridia.Clostridiales.Ruminococcaceae.Gemmiger | 3.7376 | Normal | 3.3212 | 6.46E-08 |
| Bacteria.Bacteroidetes.Bacteroidia.Bacteroidales.Paraprevotellaceae.Paraprevotella | 3.5312 | Normal | 3.2350 | 1.83E-09 |
| Bacteria.Firmicutes.Bacilli.Bacillales.Thermoactinomycetaceae.Thermoactinomyces | 1.1820 | Normal | 3.2155 | 0.000247 |
| Bacteria.Firmicutes.Erysipelotrichi.Erysipelotrichales.Erysipelotrichaceae.Allobaculum | 1.4070 | Normal | 2.9040 | 0.00122 |
| Bacteria.Firmicutes.Clostridia.Clostridiales.Clostridiaceae.Sarcina | 1.3461 | Normal | 2.9022 | 0.009918 |
| Bacteria.Firmicutes.Clostridia.Clostridiales.Lachnospiraceae.Anaerostipes | 2.9886 | Normal | 2.6412 | 9.85E-08 |
| Bacteria.Actinobacteria.Coriobacteriia.Coriobacteriales.Coriobacteriaceae.Adlercreutzia | 2.3628 | Normal | 2.5946 | 3.57E-12 |
| Bacteria.Proteobacteria.Betaproteobacteria.Burkholderiales.Comamonadaceae.Comamonas | 1.8375 | Normal | 2.5262 | 5.37E-07 |
| Bacteria.Firmicutes.Clostridia.Clostridiales.Ruminococcaceae.Butyricicoccus | 2.5880 | Normal | 2.5088 | 1.80E-08 |
| Bacteria.Firmicutes.Clostridia.Clostridiales.Clostridiaceae.02d06 | 2.1963 | Normal | 2.4993 | 0.000325 |
| Bacteria.Firmicutes.Bacilli.Turicibacterales.Turicibacteraceae.Turicibacter | 2.5207 | Normal | 2.4610 | 2.98E-07 |
| Bacteria.Firmicutes.Clostridia.Clostridiales.Veillonellaceae.Mitsuokella | 2.6093 | Normal | 2.4438 | 6.51E-11 |
| Bacteria.Firmicutes.Clostridia.Clostridiales.Clostridiaceae.SMB53 | 2.5544 | Normal | 2.3772 | 3.77E-07 |
| Bacteria.Lentisphaerae.Lentisphaeria.Victivallales.Victivallaceae.Victivallis | 2.1348 | Normal | 2.2153 | 1.28E-06 |

**Table S4 Survival state and basic information of 32 patients (paired samples)**

| SampleID | Sex | Age | Time of operation | Survival state of now | Time of death or recidivation |
| --- | --- | --- | --- | --- | --- |
| A118 | female | 69 | 2017/6/10 | survival | - |
| A122 | female | 69 | 2017/6/15 | survival | - |
| A125 | female | 56 | 2017/6/22 | survival | - |
| A143 | female | 59 | 2017/5/22 | survival | - |
| A145 | female | 73 | 2017/5/19 | survival | - |
| A011 | female | 48 | 2016/11/2 | survival | - |
| A119 | male | 62 | 2017/6/14 | death | 20180823 |
| A124 | male | 63 | 2017/6/21 | survival | - |
| A127 | male | 62 | 2017/6/22 | survival | - |
| A136 | male | 52 | 2017/5/3 | survival | - |
| A138 | female | 59 | 2017/5/4 | survival | - |
| A139 | male | 52 | 2017/5/5 | survival | - |
| A147 | male | 72 | 2017/5/24 | recidivation | 20190110 |
| A153 | male | 87 | 2017/6/9 | survival | - |
| A155 | male | 71 | 2017/6/22 | survival | - |
| A158 | male | 75 | 2017/7/12 | survival | - |
| B101 | male | 55 | 2017/7/14 | survival | - |
| B102 | female | 44 | 2017/7/12 | survival | - |
| B103 | female | 85 | 2017/7/12 | death |  |
| B104 | female | 56 | 2017/7/21 | survival | - |
| B105 | female | 35 | 2017/7/17 | survival | - |
| B109 | male | 62 | 2017/7/27 | survival | - |
| B112 | male | 67 | 2017/8/1 | death |  |
| B123 | female | 57 | 2017/8/18 | survival | - |
| B126 | male | 63 | 2017/8/25 | survival | - |
| B106 | male | 71 | 2017/7/25 | death |  |
| B114 | male | 62 | 2017/8/9 | survival | - |
| B117 | male | 68 | 2017/8/11 | survival | - |
| B122 | male | 70 | 2017/8/16 | survival | - |
| B124 | male | 74 | 2017/8/22 | survival | - |
| B127 | male | 70 | 2017/8/24 | survival | - |
| B129 | male | 66 | 2017/8/31 | survival | - |

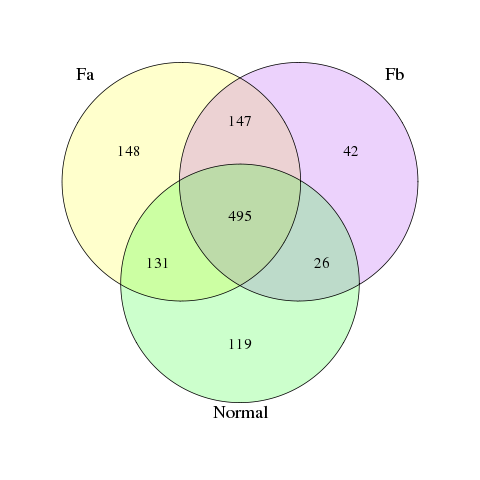

**Figure S1**. Venn of OTUs in all groups.

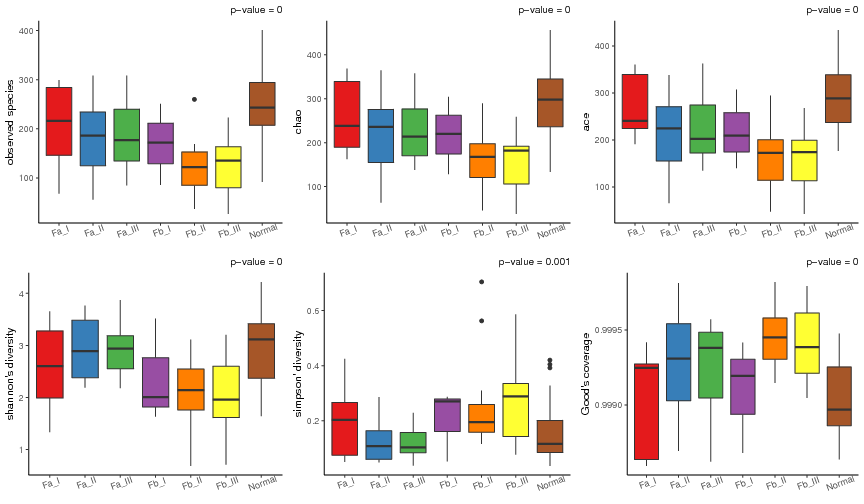

**Figure S2**. Alpha diversity. Observed species, Chao and Ace reflected community richness; Shannon and Simpson reflected community diversity; Good’s coverage reflected the sequencing coverage.

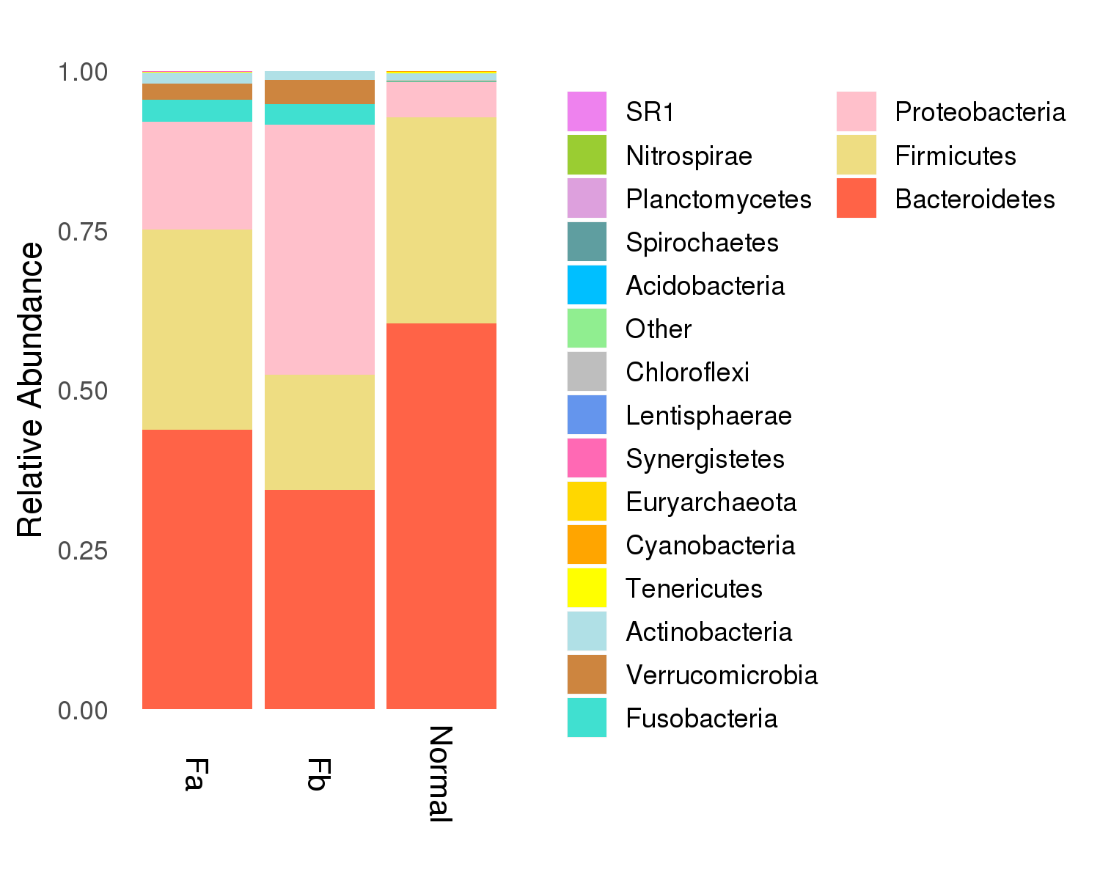

**Figure S3**. Microbiota composition of each group at the phyla level. The relative abundance less that 0.5% in all samples were combined as others.

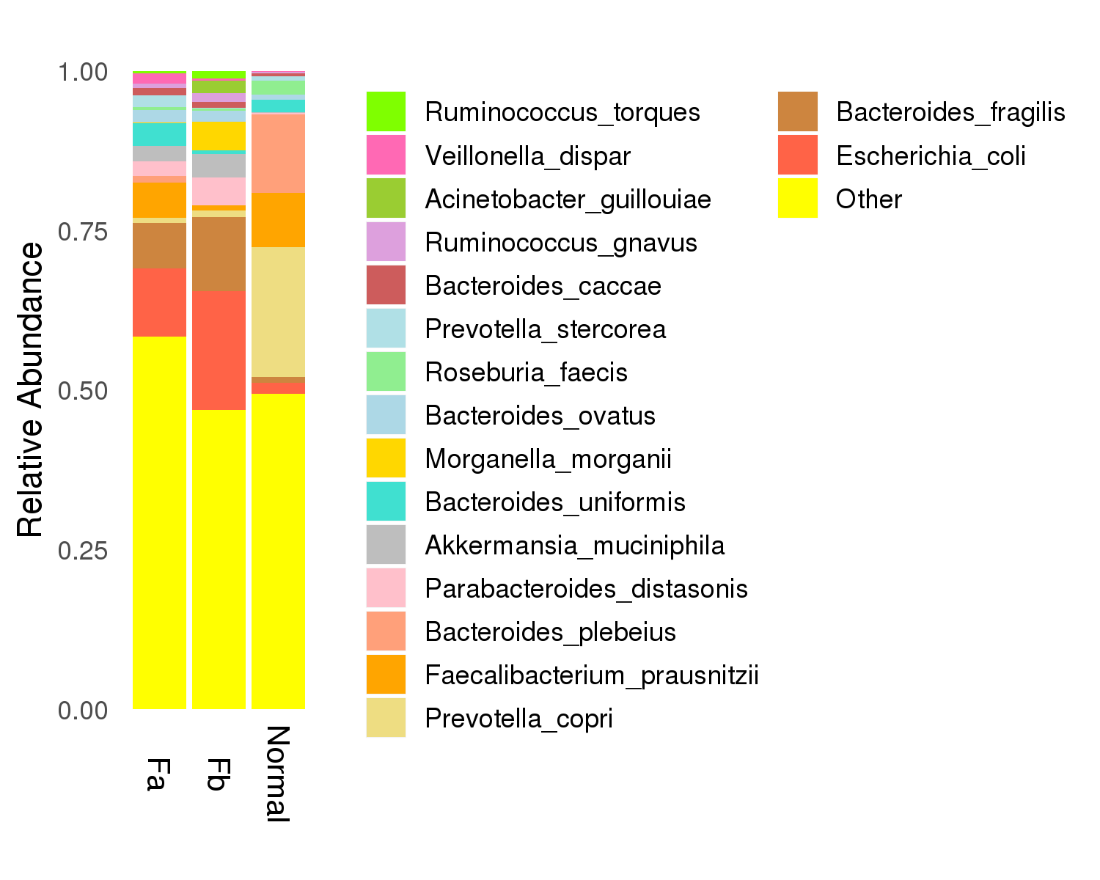

**Figure S4**. Microbiota composition of each group at the species level. The relative abundance less that 0.5% in all samples were combined as others.
